# Ultra-fast variant effect prediction using biophysical transcription factor binding models

**DOI:** 10.1101/2024.06.26.600873

**Authors:** Rezwan Hosseini, Ali Tugrul Balci, Dennis Kostka, Nathan Clark, Maria Chikina

## Abstract

Sequence variation within TF binding sites can significantly affect TF-DNA interactions, influencing gene expression and contributing to disease susceptibility or phenotypic traits. Despite recent progress in deep sequence-to-function models that predict functional output from sequence data, these methods perform inadequately on some variant effect prediction tasks, especially with common genetic variants. This limitation underscores the importance of leveraging biophysical models of TF binding to enhance interpretability of variant effect scores and facilitate mechanistic insights.

We introduce *motifDiff*, a novel computational tool designed to quantify variant effects using mono and dinucleotide position weight matrices. *motifDiff* offers several key advantages, including scalability to score millions of variants within minutes, implementation of statistically rigorous normalization strategy critical for optimal performance, and support for both dinucleotide and mononucleotide models. We demonstrate *motifDiff* ‘s efficacy by evaluating it across diverse ground truth datasets that quantify the effects of common variants in vivo, thereby establishing robust benchmarks for the predictive value of variant effect calculations. Finally, we show that our tool provides unique insights when interpreting human accelerated regions. *motifDiff* is available as a standalone Python application at https://github.com/rezwanhosseini/MotifDiff.

## Introduction

Understanding the functional consequences of genetic variants is a cornerstone of modern genomics and molecular biology. As our ability to identify genetic variation associated with disease and phenotypic traits continues to advance, so does the need for efficient tools to assess the potential impact of these variants on gene regulation. One crucial aspect of gene regulation is the binding of transcription factors (TFs) to specific DNA sequences. Variants within TF binding sites can disrupt or enhance TF-DNA interactions, thereby influencing gene expression and ultimately contributing to disease susceptibility or phenotypic traits.

In this context, a fast and reliable computational tool that can rapidly quantify the effect of genetic variants on TF binding is of critical importance. In recent years, noncoding variant interpretation has been driven by the use of deep sequence-to-function models [1, 2, 3, 4] that are trained to predict binding assay output directly from sequence. However, while powerful, these methods are not without their limitations. It has been recently shown by our group and others that they perform nearly randomly on some variant effect prediction problems when assessing human common variants[5, 6]. Without an underlying mechanistic model, this surprising lack of predictive power cannot be easily analyzed. Therefore, quantifying variant effects from biophysical models of TF binding remains an important approach. The biophysical perspective enables more limited but interpretable models, and thus can provide a critical link between molecular mechanisms and black-box predictions.

Here we address this need by introducing a highly efficient tool, *motifDiff*, designed to quantify variant effects in the context of mono and di-nucleotide position weight matrices (PWMs) that model TF-DNA interaction. *motifDiff* makes several contributions distinguishing it from previous efforts in this domain [7, 8, 9]. It is highly scalable and able to score millions of variants in minutes. It implements several normalization strategies, including an exact correction mapping motif scores to probabilities, which we demonstrate is critical for favorable performance. And it supports di-nucleotide as well as mono-nucleotide PWM models. Finally, we evaluate our method on a variety of datasets quantifying effects of common genetic variants in vivo, which establishes robust benchmarks for measuring the predictive value of variant effect calculations.

### Approach overview

#### *motifDiff* methodology

motifDiff is based on modeling the interaction of TFs with DNA via *TF motifs*. In this approach, TF motifs can be viewed as generative models for DNA sequences [10]. Sequences from a TF binding site are modeled by a position-specific probability matrix (PSPM), where element *ij* represents the probability of nucleotide *i* at motif/binding site position *j*. Combined with a background distribution (e.g., marginal nucleotide frequencies of the PSPM), the PSPM entries are transformed into log-odds by dividing each probability by its background counterpart. This results in a position-specific scoring matrix (PSSM) or position weight matrix (PWM). Rahmann et al. [10], for example, formalized this approach and provided an efficient algorithm to compute the exact distribution of log odds scores given a PSPM and background model.

A straightforward approach could be to quantify the effect of a DNA sequence variant by subtracting the log-odds/PWM score of the reference (REF) vs. the alternative (ALT) variant, or the respective maxima of variant-overlapping windows the size of the motif length. However, this approach forfeits information about the absolute values of the PWM scores of REF and ALT, and of its effect on TF binding. To illustrate, assume the probability of TF binding is related to the PWM score *s* with a monotone function *f* (*s*) with range [0, 1]. Considering Figure 1, panel (2), right side, as a hypothetical example of such a curve, and clearly PWM score differences between small scores (left side of x-axis) have less impact on TF binding probability than the same differences would have between larger scores (right side of x-axis). However, the true *f* (*s*) is not only typically unknown but also context-dependent. In particular, it depends on the concentration of the TF in question and likely on other factors, like the concentration of other TFs and cellular variables, such as post-translational modifications, that may affect TF affinity. To allow for a scalable and general approach, motifDiff used an approach we term probNorm, where we approximate the function *f* (*s*) with *g*(*s*), which is the cumulative distribution function of the PWM derived from the score distribution discussed above.

**Figure 1.**
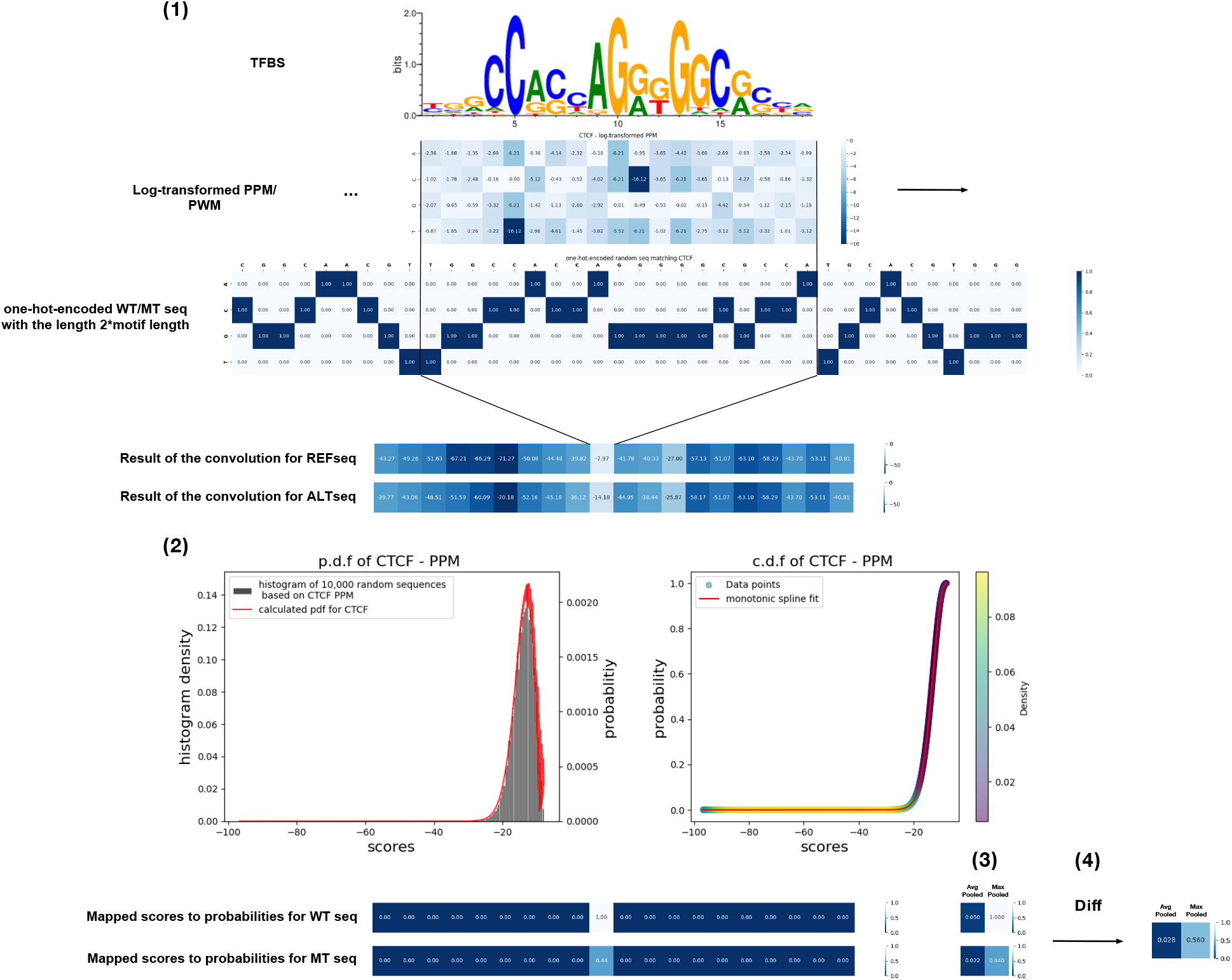
overview of the method including 4 steps: (1) mathematical convolution between the one-hot-encoded sequence (WT or MT) and the log-transformed PPM of a TFBS, (2): mapping the convolution scores to probabilities based on the motif distribution (normalization), 3: pooling (whether to choose the maximum (best match to the motif) or the average (average occupancy of the sequence in the motif distribution) of the convolution scores, and 4: getting the difference between the scores from WT and MT sequenced to see if the best match or average occupancy of the sequence is increased (constructive) or decreased (destructive) by the variant

For this newly proposed probNorm method we investigate two settings that differ by how the per-position score is summarized (max *vs*. average) across a PWM-scanned and probability-transformed DNA sequence. While taking the maximum binding position for both ALT and REF is a natural choice, the maximum match position can switch between the REF and the ALT, and thus in the max-pool setting, the REF and ALT scores would correspond to different binding events. This is naturally handled by averaging, which considers all possible binding positions. When the scores are normalized to more accurately reflect binding probability, however, the contribution from low-affinity positions is near 0, making the difference between max and average subtle. On the other hand, average pooling on raw scores has no natural interpretation and, as expected, drastically reduces performance. Consequently, we omit average pooling for un-normalized scores from our comparisons.

For TF motifs, we use 771 HOCOMOCO human PWMs for all our calculations. Our computational frameworks is flexible and allows us to test different configurations of the variant effect computation. For comparisons, we take raw log-odds difference (“No normalization”) as a baseline. We also re-implement the normalization employed in the FABIAN tool [7], which maps PWM-based scores to the [0, 1] range heuristically.

We also note that our implementation supports different window sizes (around a genetic variant) for matrix scanning. By default, the window is based on the motif length, so scores from PWM positions that do not overlap the variant are not considered. While this is a natural choice, it could be suboptimal for some downstream applications. For example, if a variant is near but not overlapping a strong TFBS it may be beneficial to consider the effect of that variant on that specific TF be 0. We find that motif-based window was optimal in our evaluations (see Supplemental Figure S1) and do not consider longer windows further. However, as the correct choice depends on the downstream application, we leave the wider-window option to the user.

#### Gold standards for variant effects

The goal for our evaluations is to closely match the typical use case for interpreting non-coding variants in the context of human phenotypes, and we compiled our validation datasets accordingly. Specifically we required that the datasets directly measured the effect of a large collection of naturally occurring variants *in vivo*. The restriction to *in vivo* assays is motivated by the need to model the complexity of TF binding mechanisms which may include interactions with co-factors and chromatin context. The focus on natural variation is likewise critical as we expect that large effects on functional binding sites are actively selected against [11, 12] and common variants are more likely to influence phenotypes through subtle and quantitative changes.

The studies that satisfy these requirement are primarily of two types. One approach is to perform an assay across a population of individuals and assess the statistical relationship between a variant and a molecular readout via quantitative trait locus (QTL) analysis. Because of linkage disequilibrium, QTL associations are not necessarily causal. Since we do not expect to be able to predict non-causal effects from biophysical models of TF binding we do not expect *a priori* that perfect predictive performance is possible. Within the caQTL dataset (see Table 1)it is possible to enrich for causal variants by looking for concordance of variant effect across all populations, and this is the finemapping approach taken by the original study. No finemaping information was available for the bQTL dataset.

**Table 1:**
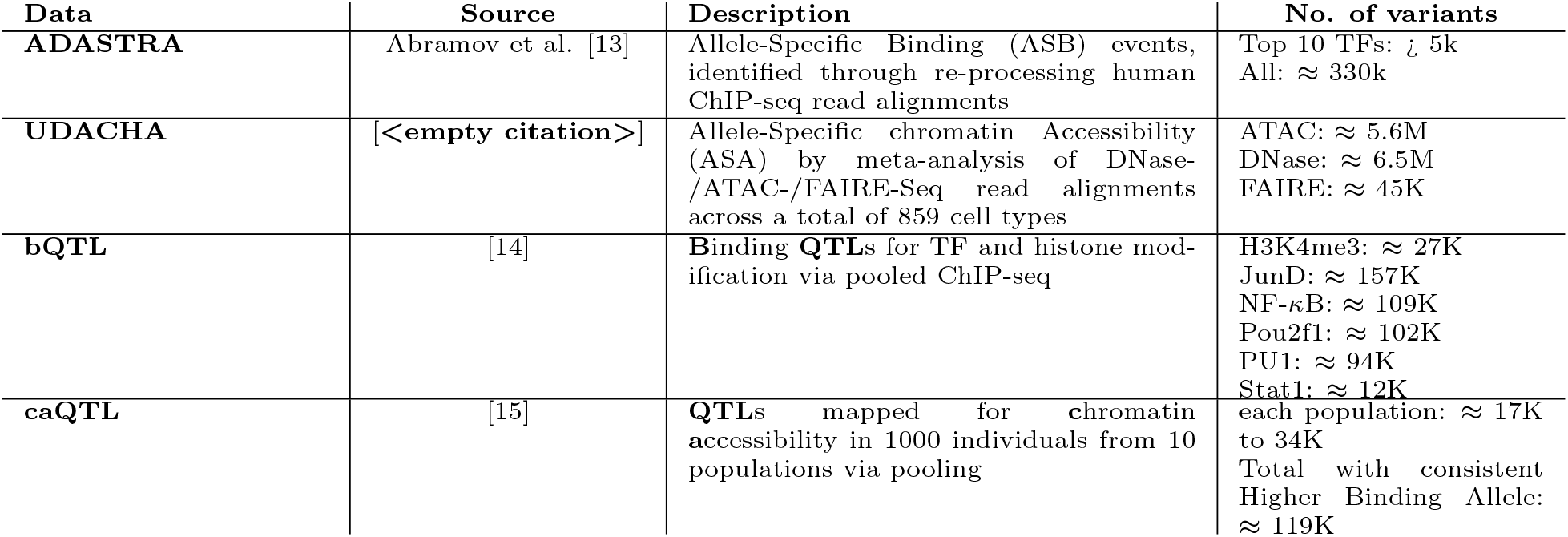
Datasets used to construct gold-standard for variant effect prediction tasks.

The second approach is to use single genotype sources but leverage heterozygosity by mapping allele-resolved reads to the correct haplotype. This approach is taken in the ADASTRA and UDACHA datasets that quantify allele specific TF binding and open chromatin respectively. It is important to note that this approach, while not sensitive to long-range LD, still is not able to distinguish between two linked alleles if they often appear in the same sequencing read. Thus, we expect this approach to produce set of alleles that is likely to be causal, but not without error.

While not all of these datasets are measuring TF binding they all measure *local chromatin features* and we make the assumption that these are themselves a function of the local binding of DNA specific proteins. This assumption is well supported by the observation that local chromatin features can be predicted from local sequence with high accuracy [1, 16, 2]. Importantly, even in the case of a specific TF binding assay, the assay output cannot be reduced to the direct affinity between the protein and the sequence. Other co-factors and general chromatin context play a role. To account for all of these effects we treat each variant effect prediction method as a feature generator that is then fine-tuned to predict the specific allelic effect task through an elastic-net regression layer trained identically for all inputs.

The fine-tuning approach allows us to investigate any dataset that reports variant effects on local chromatin features without needing to know the exact TFs involved. In the few cases where a direct comparison between a single motif score and the variant effect task is possible, we find that the ranking of the variant effect methods is consistent across elastic-net fine-tuning and single motif scores.

Finally, we note that while the predictive power varies considerably both by the variant effect method and the test dataset they are all relatively low. PWM-based methods do not achieve a correlation of above 0.5 in any setting and often the actual value is much less. This relatively low predictive performance could be attributed to a number of factors. Potential sources of low predictive power include 1) missing relevant PWMs or other sequence features, 2) sub-optimal binding approximations 3) non-linear effects not captured by linear fine-tuning and 4) noise in the test dataset. The test dataset may indeed contain significant noise due to residual confounding by linked non-causal alleles and insufficient statistical power (low minor allele frequency or few allele resolved reads). In order to distinguish between modeling limitations and noise in the test dataset we compare our variant diff scores to a state of the art deep learning approach, Sei [2]. Using Sei implicitly solves problems 1-3 as it is trained to specifically predict binding, and thus already computes the optimal convolutional feature extractions, binding approximation, and any non-linear effects. As expected, the Sei features achieve the best predictions in many (though not all) evaluations. However, the maximum correlation achieved in any setting is improved only to 0.65 indicating that the low performance is mainly a function of dataset noise rather than modeling limitations.

## Results

### *motifDiff* with probNorm is an effective approach for quantifying non-coding variant effects

We demonstrate our evaluation approach using the ADASTRA dataset, which computes allele-specific binding for transcription factors from allele-resolved reads. This dataset is significant because it directly measures TF binding (unlike other chromatin features) and is relatively well predicted. Our focus is on the 10 TFs with the most reported variant effects. We assess the holdout correlation for a linear regression task predicting the value of the variant effect (Figure 2, top panel). Since the quantification of effect size depends on statistical power, which varies by variant, we also report performance on the binary prediction task for the direction of effect (see Figure 2, lower panel).

**Figure 2.**
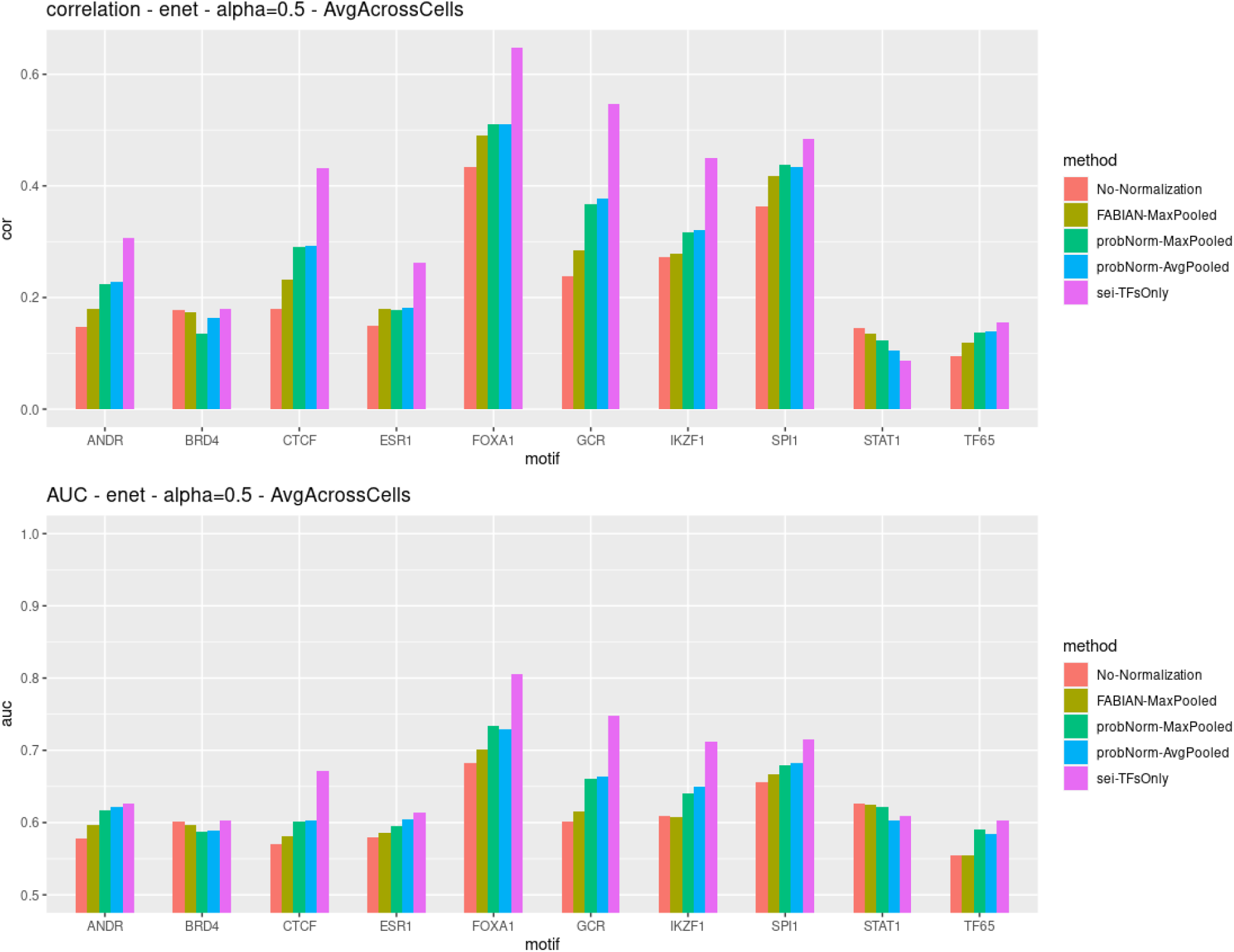
Correlation and AUROC from elastic-net prediction using scores calculated by four methods: logodds diff, FABIAN, probNorm-avgpool probNorm-maxpool for variants in the ADASTRA dataset [13]. The purple bar corresponds to a variant diff features produced by the deep learning model Sei and is included for reference as an upper bound on predictive performance. The upper panel shows correlation of predicted with observed values, lower panel shows classification performance of the observed direction of the variant effect.

As expected, we find that the variant effect features produced by the deep learning method, Sei, perform best on most ADASTRA tasks, establishing a valuable performance benchmark. Among the methods that directly use PWMs, we find that the unnormalized score difference is inferior in 8 out of 10 tasks, while the PDF normalized method we propose performs best in 8 out of 10 tasks. We observe a consistent trend where for tasks that can be well predicted (as measured by Sei’s performance), the sequence of performance is un-normalized *≤* FABIAN *≤* ProbNorm *≤* Sei indicating that increasingly sophisticated methods perform better. For the two tasks in which Sei does not emerge as the top performing method (BRD4 and STAT1), we also do not observe a consistent trend among PWM methods. In these cases it is likely that the prediction problem is not well specified as even the comprehensive Sei output does not appear to have highly informative features. We also find that the AUC performance for the sign of the variant effect tracks with the correlation performance, indicating that improvement in variant effect prediction is not concentrated in large effects.

Since the ADASTRA dataset is derived from ChIP-seq data of specific transcription factors (TFs), it enables the direct evaluation of single-feature variant effects without the need for fine-tuning. To compare the effects of single TFs against ground truth, each TF was matched to its corresponding motif using the symbol name. This matching process is streamlined by the fact that the ADASTRA dataset and the HOCOMOCO database, which provides Position Weight Matrices (PWMs), are both products of the same research group and have consistent nomenclature. Notably, when matched PWMs are available, the ADASTRA dataset offers its own PWM-based difference score, which we include in our comparison (Figure 3, grey bars). We find that all of the normalization schemes implemented in our tool improve upon the ADASTRA reported diff score, un-normalized log-odds was typically though not always worse and probNorm was best in all cases with overall good performance (exclusive of BRD4, IKZF1 and STAT1).

**Figure 3.**
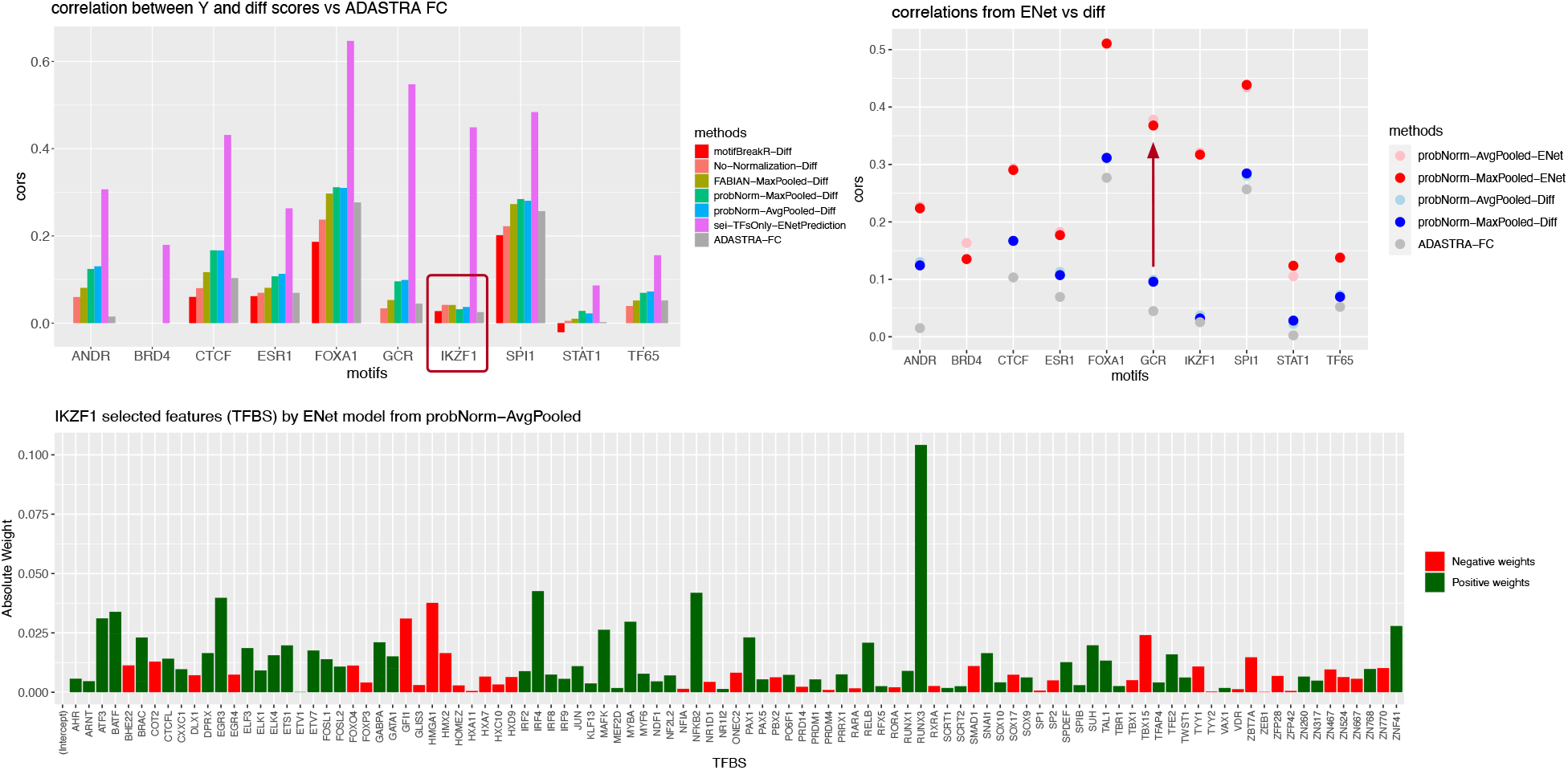
(top left) Performance of uni-variate single TF scores at predicting the affect on TF binding. Sei multi-variate predictions are included for reference. (top right) The gap between uni-variate and multivariate predictions. (bottom) TFs selected in the IKZF1 elastic-net model. Note IKZF1 itself is not included explaining the large performance gain over the univariate model

On this single score task we also compare the performance of another widely used motif scoring tool motifbreakerR [8]. This tools was used recently in several high impact studies [17, 18]. Given that this tool performed below probNorm on variant effects and was 180-400 times slower (see timing section below) we omit it from further comparisons.

Our findings indicate that, compared to elastic-net fine-tuning, using the matched motif score directly overall reduces performance, and in some cases dramaticall (e.g. IKZF1, BRD4, STAT1), suggesting that for these TFs, the PWM may not accurately represent binding affinity, or that binding partners and chromatin context play a crucial role. To understand the significant discrepancy between the elastic-net fine-tuned predictions and the single TF score, we analyzed the features selected by the IKZF1 model, see Figure 3. Our analysis revealed that a diverse set of TF features is necessary for optimal prediction of IKZF1 allele-specific binding, with the largest contribution coming from RUNX1, while the IKZF1 PWM was not selected.

This finding could be attributed to several factors. Both the PWMs and the variant effects are constructed through a computational processing of ChIP-seq data. However, PWMs are constructed from relatively few well supported binding events while variant effects are evaluated everywhere where there is sufficient haplotype resolved reads and are likely enriched for weaker binding sites or binding through co-complexes.

It is also important to consider the possibility that this observation may be influenced by bias in the alleles themselves. The alleles analyzed by ADASTRA are not random genetic variations but correspond to those naturally occurring in cell lines. Consequently, it is conceivable that alleles capable of directly altering IKZF1 binding might be negatively selected against, leaving only alleles with weaker effects that depend on interactions to be observed. This scenario aligns precisely with our research focus, as we aim to explore the impact of common variants and effects measured in vivo.

Performance on the other three gold-standard datasets is depicted in Figure 4. We observe that the pattern of performance is similar to that of ADASTRA, with the probNorm method (represented in green and blue) outperforming other methods in the PWM class on average. Additionally, we note poor performance across all methods on QTL datasets (binding bQTL or chromatin accessibility caQTL). In many instances, Sei’s performance on QTL prediction does not significantly exceed that of PWM-based methods. These results are likely due to the QTLs being a relatively smaller and noisier dataset, which makes the fine-tuning task more challenging. Given that Sei outputs more features (*≈* 4,000 vs. 771 for HOCOMOCO), the difficulty of learning the fine-tuning model is increased. Since there is no principled method to subsample Sei features to match the HOCOMOCO input dimension, a natural choice is to perform PCA; however, we found that this further decreased performance.

**Figure 4.**
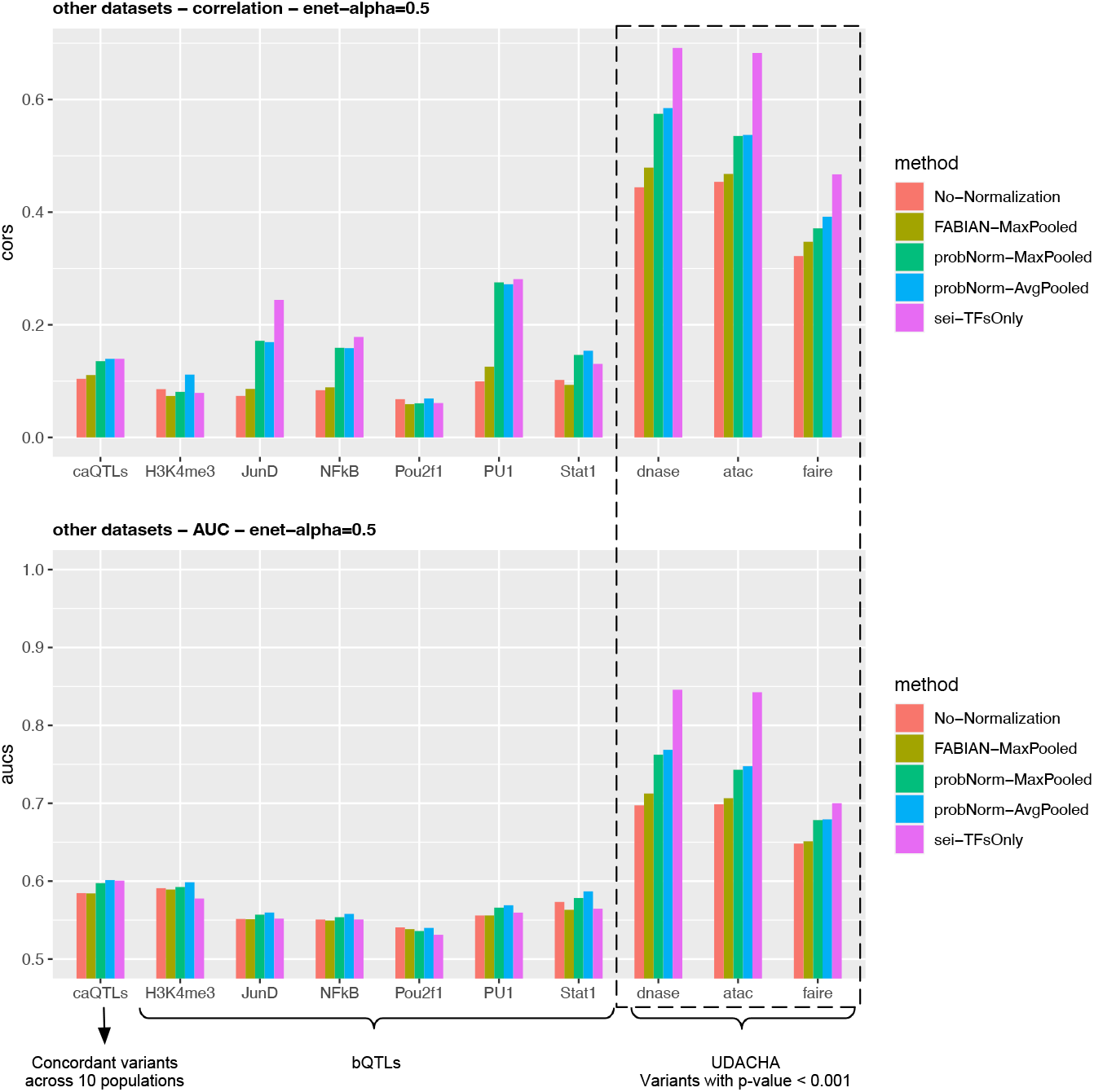
correlation and AUC from the prediction by elastic-net on the scores calculated by variant prediction methods. The results from UDACHA and two population genetic dataset are depicted on the same axis.

We note that a potential issue with the QTL datasets is that the variants may not necessarily be causal, due to LD confounding, which naturally limits the theoretically best performance that can be achieved with mechanistic models. We can investigate the contribution of LD confounding using the caQTL dataset where measurements were conducted across six different populations. We can enrich for causal variants by considering only those variants with concordant effects, as per the fine-mapping approach adopted by the original study. However, our findings indicate that performance on the caQTL dataset, even when focusing on concordant variants, does not improve (see Figure S3). This suggests that predicting QTLs is a challenging task for reasons that extend beyond the confounding effects of linkage disequilibrium (LD).

Finally, we consider the effect of max vs average pooling. Average pooling considers all possible binding positions that overlap the variant in question. Max-pooling considers the maximal site only, which has the subtle disadvantage that it may be different in the two version of the sequence being contrasted.

Average pooling resolves this issue but at an increased computational cost of multiple queries to the normalizing function. At the same time changes in the location of the maximal binding score occur rarely. Indeed, in our aggregate performance plots, probNorm-maxpool and probNorm-avepool appear similar.

However, expressing the difference as percent improvement we find that average pooling produces a small but uniform improvement across all binary prediction tasks for the variant sign (Figure 5, right panel). It also shows less consistent but still significant improvement for the quantitative effect size prediction. As the binary prediction task is independent of the magnitude and thus give more weight to small changes in binding our results suggest that averaging over all positions is indeed advantageous in this setting.

**Figure 5.**
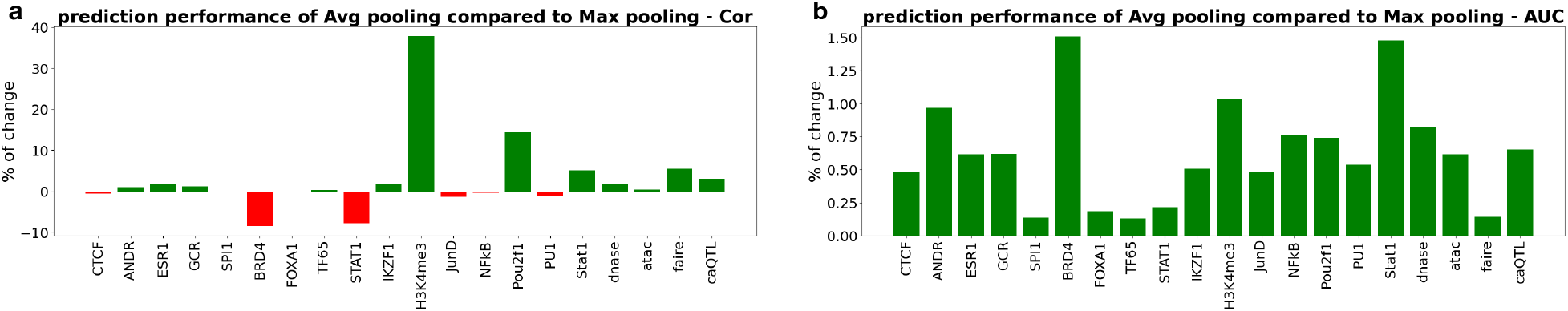
Using average pooling produces up to 40% improvement on effect sizes value prediction in some cases (a), and small but consistent improvement on all effect size sign prediction tasks (b).

### diNucleotide models do not improve variant effect prediction in the majority of cases

Position Weight Matrices (PWMs) are simplistic models based on the assumption that each base’s contribution to binding is independent. However, interdependencies between nucleotides have been consistently observed in studies [19, 20, 21, 22]. Indeed, more sophisticated models have frequently been shown to out-perform PWMs in identifying binding sites validated by experiments [23, 24, 25]. Attempts to integrate the relationships between nucleotide positions, both adjacent and distant, have led to the development of alternative approaches. Notable among these are the Binding Energy Model (BEM) [26], Dinucleotide Weight Matrices (DWMs) [27], and Transcription Factor Flexible Models (TFFMs) [23].

Our focus is particularly on DWMs because our tool utilizes fast convolutions for scalability, and DWMs can be represented as convolutions with dinucleotide-encoded sequences. Additionally, DWMs are part of the HOCOMOCO database [25] (albeit only for a subset of transcription factors), enabling a more straightforward comparison. For our evaluation, we concentrate on the common set of 278 transcription factors (TFs) represented as both DWMs and PWMs in the HOCOMOCO database. We observe that dinucleotide models generally underperform compared to mononucleotide models, with the exceptions being STAT1 and BRD4 when using the probNorm method (Figure 6). These two TFs are exceptions in that their overall performance was low, and the probNorm normalization scheme underperformed relative to no normalization. While using dinucleotide models enhances normalized performance, the non-normalized performance (illustrated by red dots in the right panel) remains comparable, suggesting that differences are likely due to interactions with normalization rather than inherent qualities of the DWMs.

**Figure 6.**
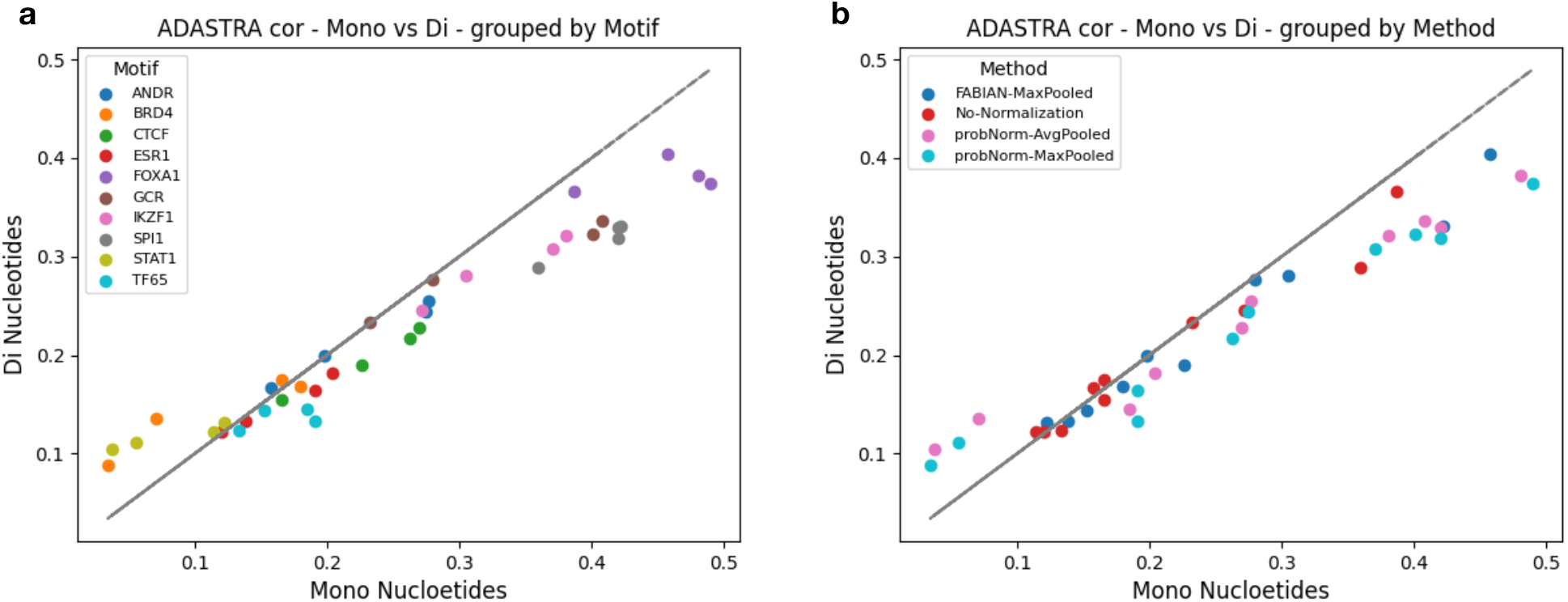
Comparing mono and di-nucleotide models on ADASTRA TF binding predictions. We find that di-nucleotide do not improve on the mon-nucleotide ones and instead suffer a decrease in performance. a, groups the performances by Motif and b, by the Method.

**Figure 7.**
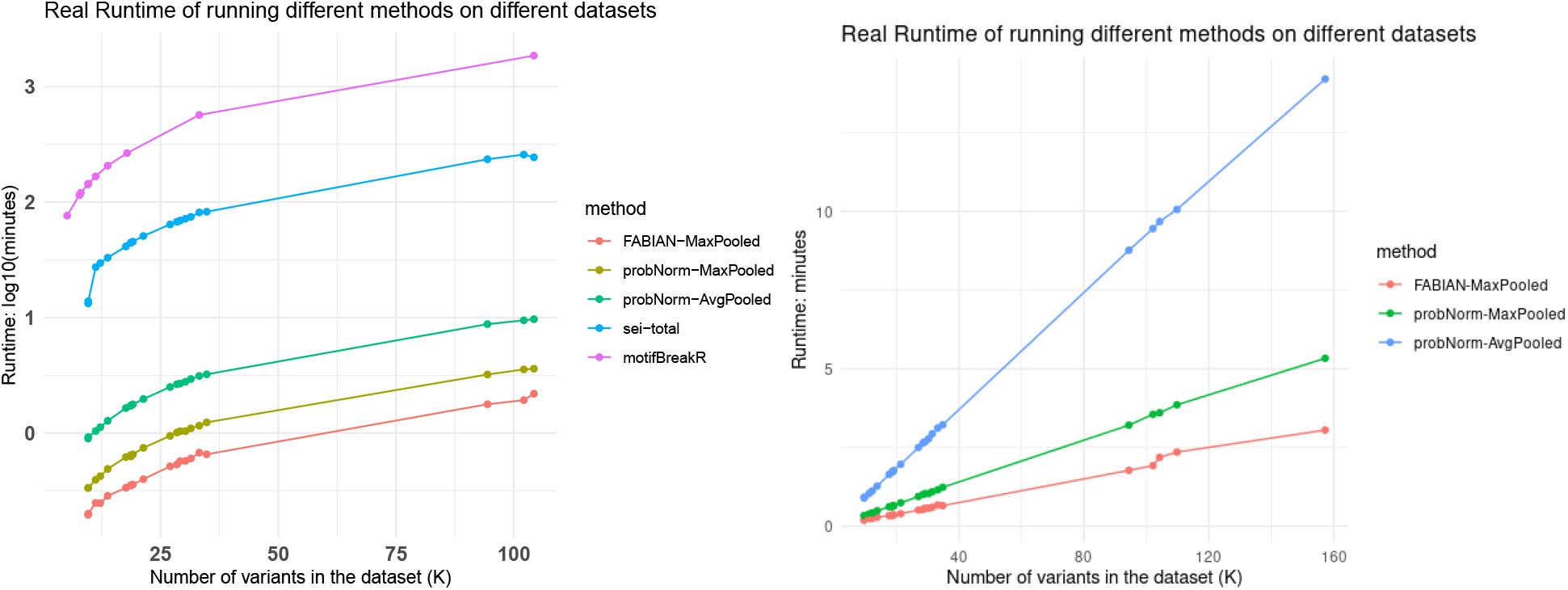
Runtimes for variant effect prediction methods plotted on the log-scale (left panel) and linear scale (right panel, excluding Sei)

In conclusion, we find no evidence that DWMs are more effective in capturing variant effects. A potential reason could be the challenge of accurately computing DWM models due to their expanded parameter space. However, the models are likely to improve as data is added and/or reprocessed and our evaluation approach can serve as a robust benchmarking platform to validate potential improvements to dinucleotide models. Additionally, the complex interplay with normalization indicates that there may be opportunities for enhancing dinucleotide performance through improved normalization strategies.

### *motifDiff* is highly scalable

One of the driving motivations for our approach is the scalability. We compare the scalability of our approach to another popular motif based method, motifbreakeR [8] which has been used in several high impact studies [17, 18]. We find that our method is more than 180-480 times faster (depending if max or average mode is considered) than motifbreakR running single threaded. We note that the probNorm normalization indeed incurs an extra cost due to queries to the empirical CDF function and this is magnified for average pooling approach. We also note that probNorm is considerably faster than Sei with a 25 to 70 fold speed up depending on the setting.

### *motifDiff* scores provide interpretable signals of TFBS change in evolutionary contexts

Evolutionary analysis is one research area where the importance of incorporating biophysical models is particularly evident. Deep learning models often inherit biases present in the genomes they are trained on. A well-documented example is the tendency of neural networks to rely on lineage-specific repetitive elements [28].

In contrast, the biochemical binding preferences of orthologous transcription factors are often deeply conserved [29], suggesting that many position weight matrices (PWMs) remain informative even in highly divergent genomes.

To investigate an evolutionary setting where the changes in TFBS are of interest, we consider the human accelerated regions (HARs) whose functionality has been extensively studied. The current view of HARs is that many of them represent neurodevelopmental enhancers that have rapidly changed their function in the humane genome and thus may underlie human specific traits [30].

We analyze the HAR variants reported in Whalen et al. [31] form the perspective of TFBS gains and losses using different approaches. The original study did not focus on TFBS level analysis opting to use the higher level Sei sequence classes. Sei sequence classes, which compresses all epigenetic tracks predictions into 40 functional categories, including promoters and different enhancer types. Using Sei functional classes, the authors found that the HARs sequences were equally likely to experience gains and losses [31].

Here we perform the same analysis from the TF perspective. Specifically, we use different TF scoring methods and compute the Empirical Probability of Positive Shift (EPPS), defined as the proportion of variants with a positive Diff score. Statistical significance was assessed using a Wilcoxon rank-sum test against a normal distribution. We visualized EPPS against − log_10_(FDR-Pvalue). This analysis revealed a overall bias toward gains (Figure 8a,b), and the transcription factors associated with gains were strongly enriched for neurodevelopmental roles.

**Figure 8.**
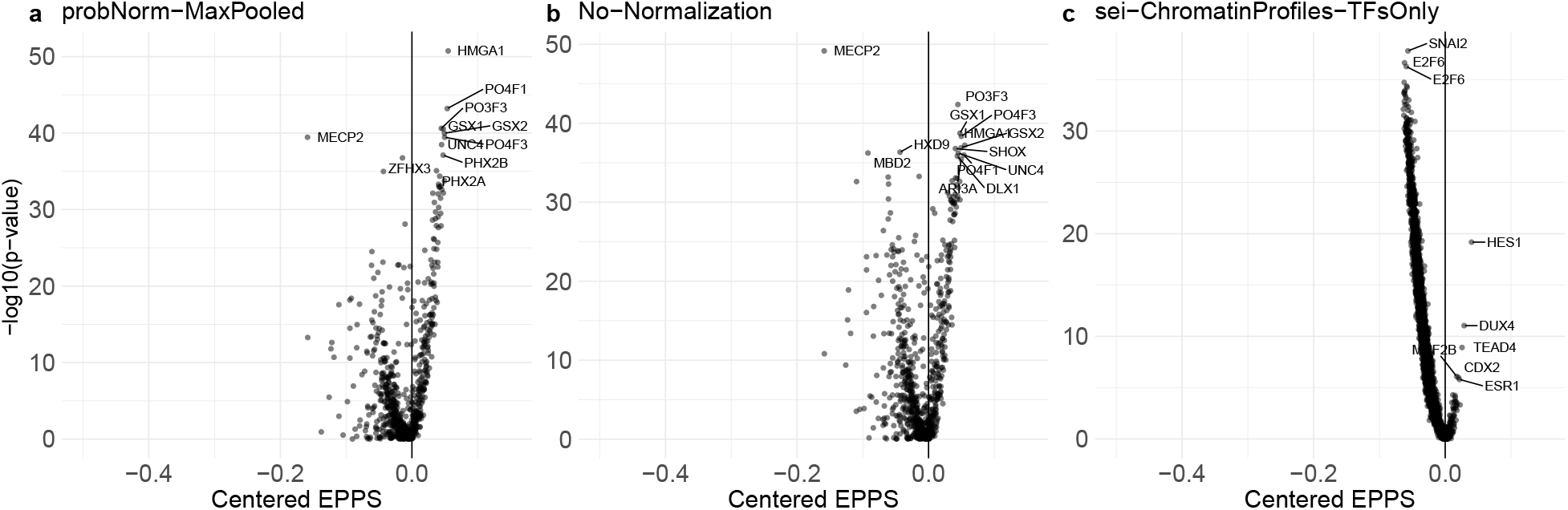
probNorm Diff scores show more stronger signals compared to Diff scores from sei. **a,b,c**. show the Positive Distribution shift from Normal Distribution on Diff scores for each TF. Since under the null hypothesis the EPPS equals 0.5 we centered it by subtracting 0.5 to clearly indicate binding gains as positive values. (a) probNorm normalized with Max Pooling (b) PWM scores with no normalization, (c) Sei transcription factor predictions.

The POU and GSX family transcription factors play key roles in nervous system development. POU3F3 (also known as BRN1) regulates cortical neuron migration and layer formation during brain development [32]. POU4F1 (BRN3A) and POU4F3 (BRN3C) are critical for the survival and differentiation of sensory neurons, including auditory hair cells, with POU4F3 specifically required for cochlear hair cell maintenance [33, 34]. GSX1 and GSX2 orchestrate regional patterning of the ventral forebrain and spinal cord [35, 36].

We also note that the enrichment in gains is more evident for probNorm when compared to no normalization (Figure 8 a and b). At the same time Sei TF track predictions (Figure 8 c) are highly enriched for losses. The gained TFs also have less direct neuronal relevance though we note that HES1, DUX4, CDX2, TEAD4 have critical roles in early development broadly including neurogensis [37, 38, 39, 40].

## Discussion

We introduce *motifDiff*, a novel tool designed for scoring variant effects using precomputed mono- and dinucleotide models of transcription factor (TF) binding preferences. We assess its performance across a variety of settings, focusing on ground truth variant effects measured in vivo on common human variants—a scenario that closely mirrors the practical application of variant prediction. This contribution is vital, as the accuracy of variant prediction methods can vary significantly depending on the context. For instance, while the complex and high-performing deep learning model Enformer delivers accurate, state-of-the-art predictions on Massively Parallel Reporter Assays (MPRA) data, it demonstrates nearly random accuracy in predicting the direction of common variant expression quantitative trait loci (eQTLs).

Moreover, our tool leverages fast convolutions for sequence scanning, enabling us to conduct comprehensive experiments across different settings. We discover that, as anticipated, raw log-odds differences are not the most effective measure due to the non-linear relationship between log-odds and binding probabilities. While heuristic methods like those employed by FABIAN offer improvements over raw log-odds, our proposed “probNorm” approach is both more principled and more performant.

As part of our evaluation, we also explore how *motifDiff* can reveal biologically meaningful patterns in an evolutionary context. Using human accelerated regions (HARs) as a case study, we show that *motifDiff* highlights consistent TFBS gains and losses associated with neurodevelopmental regulatory changes. This analysis demonstrates how even simple, interpretable biophysical models can capture evolutionary signatures that may be obscured in more complex frameworks.

In summary, *motifDiff* offers a scalable and refined method for scoring variant effects through biophysical models of TF binding. Although simple in design, this mechanistic framework serves as a foundational element that can be integrated into more complex models, as demonstrated by our application of linear fine-tuning for tasks downstream of TF binding, such as identifying open chromatin regions. Additionally, we envision that this TF-centric framework will offer valuable insights for interpreting both the successes and limitations of powerful black-box models.

