## Supplementary data for "Ultra-fast variant effect prediction using biophysical transcription factor binding models"

### Supplementary Methods and Figures

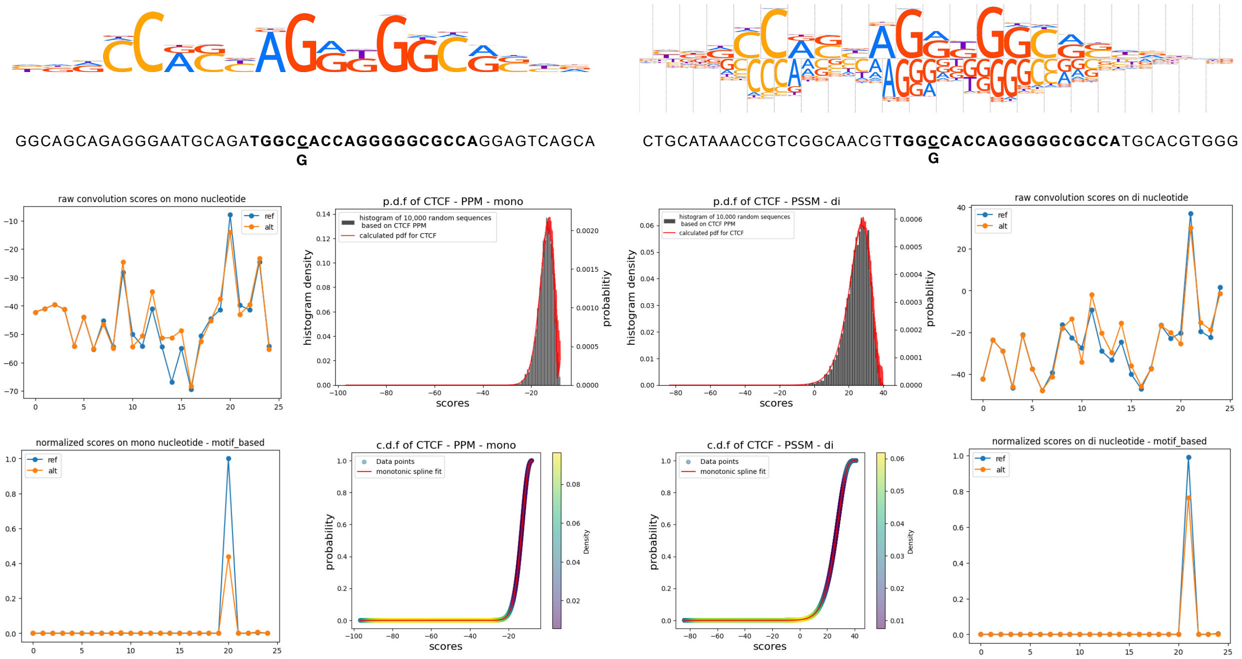


Supplemental Figure S1: Affects of the normalization on a single variant affecting a high scoring CTCF site for both mono (left) and di (right) models. Only one of the possible positions has high binding affinity and this position corresponds to the max score. However, in terms of the difference between REF allele in yellow and ALT allele in blue there are many ”non-binding” positions that generate comparable or greater difference in scores illustrating why raw score difference is not optimal for variant effect quantification. On the other hand after the probability normalization only the binding positions have scores that are appreciably different from 0. We also note that normalization is equally effective for the dinucleotide model

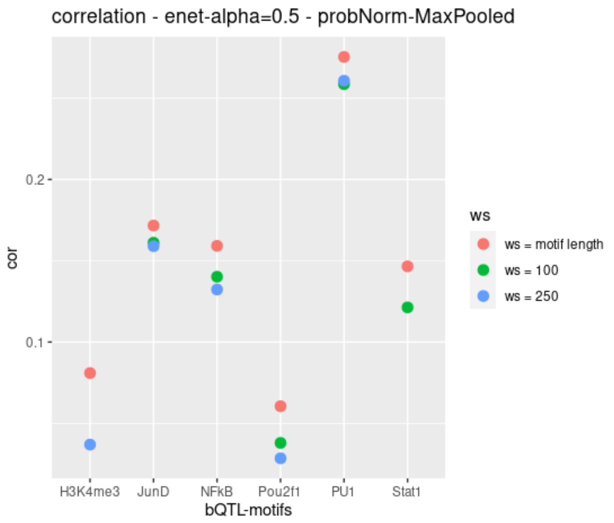


Supplemental Figure S2: Effects of changing the window size on bQTL predictions. Since bQTL data only reports an association of a variant with a binding peak it is possible that we can improve the performance by considering a larger window around the variant when evaluating the initial PWM scan. In this setting, a variant that is near but not strictly inside a strong binding site will not affect the maximum score and thus the variant effect will be 0. On the other hand, when only variant overlapping positions are considered the variant effect is always different from 0 though it can be very close to 0 when using FABIAN or probNorm normalization. We find that in our evaluations increasing the window size reduced performance.

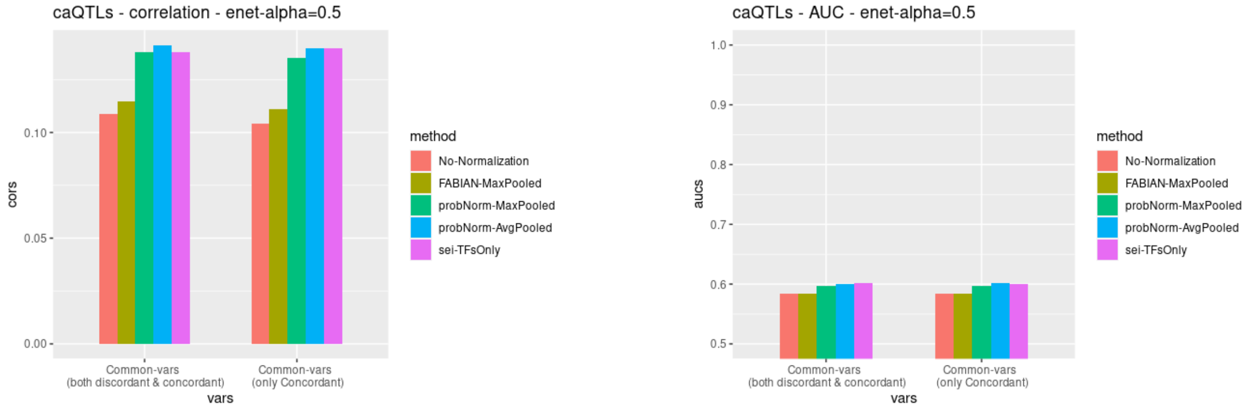


Supplemental Figure S3: Comparing the predictions for all chromatin accessibility (caQTL) variants averaged across populations against the concordant subset. The concordant ones are expected to be enriched for causal variants as LD structure varies across populations. We fine that focusing on concordant variants did not appreciably improve predictions

### Detailed Methods

The method we present in this paper is to calculate a score, quantifying the effect of non-coding variants on transcription factors binding to the site. The score is calculated through multiple steps including: 1) the mathematical convolution between a selected sequence, with reference or alternative allele (REF, ALT), and the Position Probability Matrix (PPM) of a TFBS, 2) mapping the scores from the convolution to probabilities, 3) pooling all the scores from each possible position into a single score, and 4) computing the difference between the scores from REF and ALT.

Given a variant and a PWM we score the PWM log-odds within a given window of the variant. By

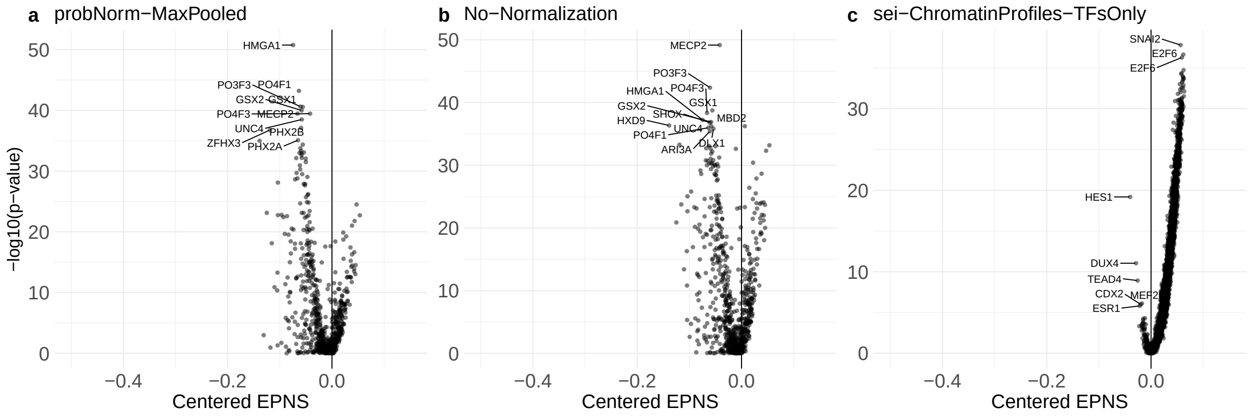


Supplemental Figure S4: Diff score Negative Distribution shift from Normal Distribution. We centered the EPNS at 0.5 to show the TFs with bindings losses in positive values. The TFs with binding losses are more clearly recognized by probNorm Diff scores, while in the No-Normalization, the raw log-odd scores are not able to effectively differentiate between the TFs with binding losses and the others.

default the window is twice the length of the motif, so scores from PWM positions do not overlap the variant are not considered. While this is a natural choice it could be suboptimal for some downstream applications. For example if a variant is near but not overlapping a strong TFBS we may be beneficial to consider the effect of that variant on that specific TF be 0. As the correct choice depends on the downstream application we leave the wider-window option to the user. The selected sequence is then converted to a one-hot-encoded array, to go through a discrete convolution with the log-transformed PPMs considering both forward and reverse complements given the final log-odds score. Here we are using the 771 human TFBSs provided by HOCOMOCOv11 [1] (for a t total of 1542 convolutional filters).

Sliding the selected sequence of length ls, over the motif of length $l_m$ by nucleotide steps and calculating the convolution at each step, we get $l_s - l_m + 1$ scores, each representing a measure of how similar the sequence at that position is to the motif. After all the scores are mapped to probabilities we apply either average or max pooling on them to extract only one value for the variant in each sequence and each TFBS. The maximum value represents the probability of the best possible match between the sequence and the motif and the average value represents the average occupancy of the motif on the selected sequence.

#### Normalization

Each motif is represented by a Position Probability Matrix, $M \in R^{l \times n}$ where columns correspond to single nucleotides, if n=4, or dinucleotides, if n=16, in the lexicographical order and rows to motif positions. Considering a background probability for single nucleotides, $\pi = \pi(\pi_A, \pi_C, \pi_G, \pi_T)^T$, we can get the probability of a nucleotide sequence of length $l(x_1|x_2|...|x_l)$ as:

$$P((x_1|x_2|...|x_l)) = \prod_{i=1}^{l} \pi(x_i)$$

Where $x_i$ is the nucleotide at $i$ th position and could be A, C, G, or T. the motif score for the sequence is defined as:

$$score((x_1|x_2|...|x_l)) = \sum_{i=1}^{l} M[i, x_i]$$

if considering single nucleotides, and:

$$score((x_1|x_2|...|x_l)) = \sum_{i=1}^{l-1} M[i, (x_i|x_{i+1})]$$

if considering di-nucleotides. The motif score distribution is then calculated to be the district p.d.f of motif scores for sequences of i.i.d nucleotides:

$$S(l; x_l) = P(score(x_1|x_2|...|x_{l-1})|x_l \approx \pi)$$

Where $S(l; x_l)$ is the score distribution of sequence of length $l$, ending in $x_l \in \{A, C, G, T\}$.

**Mono-nucleotide distribution:**

When using mono nucleotide PPMs, every single nucleotide takes part in the score individually, so for the start we would only have a single nucleotide $x_i$ with no condition, with a score distribution as $S(1, x_i)$ where $x_i$ is the single nucleotide in the set {A, C, G, T}.

$$P(score(x_i)|x_i \approx \pi) = S(1; x_i) = \delta(M[1, x_i], \pi(x_i))$$

for $x_i \in \{A, C, G, T\}$, where $i$ is the position of the nucleotide in the motif (rows of PPM).

Considering $S(1; x_1)$ and $S(1; x_2)$, as the score distribution of the single nucleotide at the 1st and 2nd position respectively, we can make the score distribution for a sequences of length two by convolving $S(1; x_1)$ and $S(1; x_2)$:

$$S(2; x_2) = P(score(x_1)|x_2 \approx \pi) = P(score(x_1) \approx \pi) * P(score(x_2) \approx \pi)$$

$$S(2; x_2) = S(1; x_1) * S(1; x_2)$$

$$S(2; x_2) = \delta(M[1, x_1], \pi(x_1)) * \delta(M[2, x_2], \pi(x_2))$$

The result, $S(2; x_2)$ will be representing the score distribution of a sequence of length two. We can update the score distribution for every new nucleotide added to the sequence at each step, by convolving the score distribution of the new nucleotide with the score distribution of the sequence from the previous step:

$$S(n+1; x_{n+1}) = S(n; x_n) * S(1; x_{n+1})$$

$$S(n+1; x_{n+1}) = S(n; x_n) * \delta(M[n+1, x_{n+1}], \pi(x_{n+1}))$$

Where $\delta(x; y)$ is an "un-normalized" p.d.f with mass $x$ and value $y$. Note that since $\delta(x; y)$ only has one value, convolution of a discrete p.d.f with $\delta(x; y)$ only consists of (i) adding x to all values and (ii) multiplying with y. The final distribution $S(l; x_l)$ will be an "un-normalized" p.d.f distribution of the scores for a sequence of length $l$.

**Di-nucleotide distribution:**

When using PPMs with dinucleotides, we need to calculate the score distribution by treating the (sub)sequences that end in each of the four different nucleotides separately. We initialize the distribution for all the sequences of length two that end in A,C,G, or T, respectively as:

$$P(score(*|A)) = \begin{pmatrix} M[1, (AA)]|\pi(A)\pi(A) \\ M[1, (CA)]|\pi(C)\pi(A) \\ M[1, (GA)]|\pi(G)\pi(A) \\ M[1, (TA)]|\pi(T)\pi(A) \end{pmatrix} = S(2; A)$$

Where we assume that all nucleotides ending in A have distinct motif scores. Generalization to other ending nucleotides than A is straight forward. So, at the end of initialization we have $S(2; x_i)$ for $x_i \in \{A, C, G, T\}$. $S(2; x_i)$ is the joint distribution of two events: (a) observing a certain score for dinucleotide and (b) the second nucleotide being $x_i$. Then, we update the distribution for $n+1$th step from $n$th step as:

$$\begin{aligned} S(n+1; x_i) = \quad & S(n; A) * \delta(M[n, (A|x_i)], \pi(x_i)) + \\ & S(n; C) * \delta(M[n, (C|x_i)], \pi(x_i)) + \\ & S(n; G) * \delta(M[n, (G|x_i)], \pi(x_i)) + \\ & S(n; T) * \delta(M[n, (T|x_i)], \pi(x_i)) \end{aligned}$$

Where * denotes convolution and $\delta(x; y)$ is an "un-normalized" p.d.f with mass $x$ and value $y$. Note that since $\delta(x; y)$ only has one value, convolution of a discrete p.d.f with $\delta(x; y)$ only consists of (i) adding $x$ to all values and (ii) multiplying with $y$. After calculating the last set of joint distributions, the score distribution is the marginal distribution over final nucleotides:

$$\begin{aligned} P(score((x_1|...|x_l)) = y) = \quad & P_A(score((x_1|...|x_l)) = y) + \\ & P_C(score((x_1|...|x_l)) = y) + \\ & P_G(score((x_1|...|x_l)) = y) + \\ & P_T(score((x_1|...|x_l)) = y) \end{aligned}$$

Where the "un-normalized" p.d.f $P_A$ is given by $S(n; A)$ for $n = l$, and likewise for other nucleotides.

**Fitting a function to the CDF:**

The probability density function made from PPM is discrete and can't be used to map all ranges of scores to probabilities, so we make the C.D.F of the distribution for each motif and fit a function to it to map the scores calculated for each sequence from the convolutions to probabilities. A better fit to C.D.F results in a more accurate mapping and consequently better and more predictive scores. So the best choice would be interpolation but with low computational cost. The PchipInterpolator function from scipy python library uses monotonic cubic spline for the interpolation through an incredibly fast computation. The fitted functions and parameters are all saved for each motif in HOCOMOCOv11 TFBSs, and provided along with the code for even faster implementation.

#### Evaluation

##### Data

For evaluating our method we relied on several different datasets that measure the effects of common variants in-vivo. ADASTRA[2] (Allelic Dosage-corrected Allele-Specific human TRAnscription factor binding site) provides more than 200k allele-specific binding (ASB) events, identified through processing human ChIP-seq read alignments from GTRD database, in a framework which accounts for the allelic dosage of aneuploidy and CNVs, and read mapping bias. All the variants are provided with Benjamini–Hochberg (FDR) adjusted P-value less than 0.05, across 674 TFs (including epigenetic factors) and 337 cell types. The minimum of the FDR-correced P-value of for alt and ref alleles is considered as the p-value for the existence of an ASB effect. Because the number of allele resolved reads may be low and the effect size estimate may have large confidence intervals we use signed sign $e \log_1 0p$ as a z-score like quantity that captures both effect size and the confidence. Variants are averaged across cell-types into a single score. UDACHA [3] with a different framework, though inherited from ADASTRA, but significantly adapted to account for increased variability of read counts in chromatin accessibility data, provides more than 12 million allele-specific chromatin accessibility (ASA) events by meta-analysis of DNase-/ATAC-/FAIRE-Seq read alignments of GTRD across a total of 859 cell types. They also provide FDR-corrected P-values for each allele, minimum of which is considered as the P-value of the ASA effect (should we provide a supp. Table or just reference to their work?). We also make use of two population genetic datasets. Binding QTLs (bQTLs) provided by [4] includes thousands of QTLs specifically mapped for TF binding and histone modification by performing ChIP-seq in a pooling process for five TFs critical for immune cell development and function (NF-$\kappa$B, PU.1/Spi1, Stat1, JunD, and Pou2f1/Oct1) and one histone modification with active transcription (H3K4me3). They compare the allele frequencies in each SNP before and after ChIP-seq process per TF and they consider the ones with a significant shift in their frequency (assessed by a modified binomial test that accounts for uncertainty in allele frequency estimates), as bQTLs. Chromatin accessibility QTLs (caQTLs) are produced in a similar manner with the addition of repeated measurements in six different populations [5]. The study authors reasoned that causal variants will be concordant across populations while LD confounded non-causal variants will not be due to population-specific LD structure.

##### EPPS, EPNS

The Empirical Probability of Positive Shift and the Empirical Probability of Negative Shift for each TF are calculate as:

$$\text{EPPS}_{TF} = \frac{1}{N}\sum_{i}^{N} \text{Diff}_{TF} > 0,$$

$$\text{EPNS}_{TF} = \frac{1}{N}\sum_{i}^{N} \text{Diff}_{TF} < 0,$$

where $N$ is the number of variants and Diff is the Diff scores calculated by the method (either probNorm-MaxPooled or No-normalization) for the TF.

##### Fine-tuning for specific tasks

Our evaluation is based on using variant effects as features which are then fine-tuned to specific prediction tasks. This enables a highly flexible framework that can evaluate any measured variant effect even if the measurement itself is not TF binding (e.g. histone modification, open chromatin). We used `cv.glmnet` function in the `glmnet` R package, to predict both the direction (*sign*) and the target value. The `cv.glmnet` function uses the path algorithm to perform calculations for a series of different regularization coefficients, `lambda`, and reports hold out performance across cross validation folds. We set the number of CV folds to 10 and and the number of lambdas to 100, respectively. The elastic-net model has another hyperparamter `alpha`, which trades off contributions of L1 and L2 penalties. We found that the performance of the model with alphas 0.1, 0.5, 0.9 are almost identical, and set `alpha`=0.5 for all experiments.

The prediction target is standerdized to have mean 0 and variance 1 so that R-squared calculated as $1 - \min(cvm)$, where $cvm$ is considered to be mean cross-validated MSE for Gaussian regression. We report correlation as the square root of R-squared. For the binary prediction task of predicting the variant sign we set family="binomial" and type.measure="auc" and report the AUC is given by $\max(cvm)$ which in this setting is the mean cross-validated area under the ROC curve for the best `lambda`.

We also considered a nonlinear model as the fine-tuning layer Using xgboost R package and extensive hyper-parameter sweeps we did not observe any improvement over the elastic-net performance.
